# Rapid quantitative pharmacodynamic imaging with Bayesian estimation

**DOI:** 10.1101/017921

**Authors:** Jonathan M. Koller, M. Jonathan Vachon, G. Larry Bretthorst, Kevin J. Black

## Abstract

We recently described rapid quantitative pharmacodynamic imaging, a novel method for estimating sensitivity of a biological system to a drug. We tested its accuracy in simulated biological signals with varying receptor sensitivity and varying levels of random noise, and presented initial proof-of-concept data from functional MRI (fMRI) studies in primate brain. However, the initial simulation testing used a simple iterative approach to estimate pharmacokinetic-pharmacodynamic (PKPD) parameters, an approach that was computationally efficient but returned parameters only from a small, discrete set of values chosen *a priori*.

Here we revisit the simulation testing using a Bayesian method to estimate the PKPD parameters. This improved accuracy compared to our previous method, and noise without intentional signal was never interpreted as signal. We also reanalyze the fMRI proof-of-concept data. The success with the simulated data, and with the limited fMRI data, is a necessary first step toward further testing of rapid quantitative pharmacodynamic imaging.

## Introduction

Measuring the sensitivity of an organ to a drug *in vivo* is a common, important research goal. The traditional approach is to independently measure biological responses to a range of different doses of drug. We recently described a novel method, rapid quantitative pharmacodynamic imaging (or QuanDyn™), for estimating sensitivity of a biological system to a drug in a single measurement session using repeated small doses of drug (Black et al., 2013). In that report, we tested QuanDyn™’s accuracy in simulated data with varying receptor sensitivity and varying levels of random noise. The initial simulation testing used a simple iterative approach to estimate pharmacokinetic-pharmacodynamic (PKPD) parameters including *EC*_50_, the plasma concentration of drug that produces half the maximum possible effect *E_max_*. The iterative approach was computationally efficient but could only select *EC*_50_ from a short list of parameter values chosen *a priori*.

Here we revisit the simulation testing using a Bayesian method to provide continuous estimates of the PKPD parameters. The Bayesian approach also identifies data too noisy to produce meaningful parameter estimates (using a model selection package described below). Bayesian methods have been used successfully in other PKPD analyses (Lavielle, 2014, to cite but one example). For the present purpose we applied a Bayesian data analysis package specifically designed for efficient voxelwise analysis of 4-dimensional imaging data (Bretthorst, 2014; Bretthorst and Marutyan, 2014).

## Methods

### Simulated data

We used a standard sigmoid PKPD model (Holford and Sheiner, 1982) to create 6 time-effect curves that could reasonably represent biological signal from a pharmacological challenge study: one with no response to drug (*E_max_* = 0) and five with varying sensitivities to drug: *E_max_* = 10 and *EC*_50_ ∈ 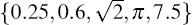.

As in the previous work, the concentration of drug in plasma over time is modeled as

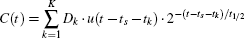

where *K* doses of drug, *D_k_*, are given at times *t_k_*, *u*(*t*) is the unit step function, *t_s_* (for “time shift”) is a fixed delay between drug concentration and effect, and *t*_1/2_ is the elimination half-life of drug from plasma (Black et al., 2013). Drug effect is modeled as

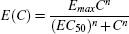

where *C* is *C*(*t*) from the previous equation and *n* represents the Hill coefficient. Baseline nonquantitative signal drift was simulated by adding to each curve a quadratic function of time

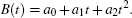

The full model is then

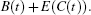

The test curves were generated using *K* = 4, *D*_1_ = *D*_2_ = *D*_3_ = *D*_4_ = the dose of drug that produces a peak plasma concentration of 1 (arbitrary concentration units), *t_s_* = 0.5 min, *t*_1/2_ = 41 min, *n* = 1, *a*_0_ = 1000, *a*_1_ = 2/(40 min), and *a*_2_ = 0. The 6 resulting curves are shown in Figure 1.

**Figure 1.**
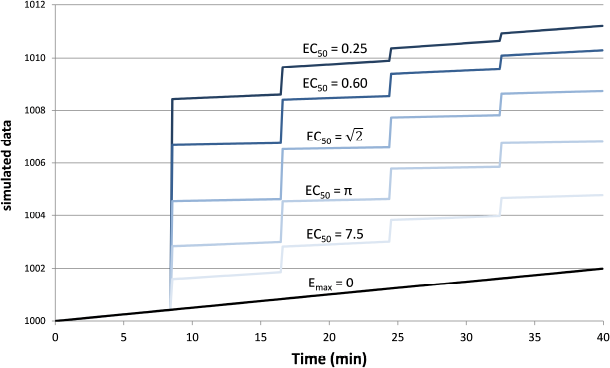
Simulated tissue responses for various values of *EC*_50_, *i.e*. the test data before adding noise.

Finally we added Gaussian noise to each time point. This was done 1000 times for each of the 6 curves above and for each of 8 noise levels from *SD* = 0.01*E_max_* to 2*E_max_*, resulting in 48,000 noisy time–signal curves plus the original 6 “clean” curves (see Supplemental Data).

### Testing the method using the simulated data

In the simulated data described above, each of the 48,006 time courses were analyzed using the “Image Model Selection” package from the Bayesian Data-Analysis Toolbox (Bretthorst and Marutyan, 2014; Bretthorst, 2014). The Toolbox computes the posterior probability for the set of models (Bretthorst, 1988) given a 4d data set. A Markov chain (Gilks et al., 1996) is used to draw samples from the joint posterior probability for all of the parameters including the choice of model. The Markov chain Monte Carlo simulation included the full model *B*(*t*)+*E*(*C*(*t*)), the baseline model *B*(*t*), and a “no signal” model. Each model has equal prior probability, or more precisely we specify that the conditional probability of any model, given the supplied prior probabilities for the parameters relevant to that model, is equal to that of any other model (see Bretthorst, 2014, section 22.1, at equation 22.6). Monte Carlo integration is then used to obtain samples from the posterior probability for each model and from the posterior probability for each parameter given the model. For the present analysis we specified 2500 samples at each step (50 samples run in parallel, repeated 50 times). Simulated annealing is used to minimize the risk of convergence to a non-global local maximum (see Bretthorst, 2014, appendix B, for details). If the posterior probability for the model indicated the full model, *B*(*t*) +*E*(*C*(*t*)), was preferred, the package also returned values for *EC*_50_, *t_s_*, *E_max_*, *a*_0_, *a*_1_, and *a*_2_. The software returns both the mean parameter values and the values from the simulation with maximum likelihood; the present report uses the latter. This analysis was repeated for each of the 48,006 time courses.

To provide more even sampling of parameter space across the conventional logarithmic abscissa for concentration-effect curves, *EC*_50_ was coded as 10*^q^*, where *q*= log_10_*EC*_50_, and a uniform prior probability was assumed for *q* with range [–3,1.3], corresponding to a wide range of *EC*_50_ values from 0.001 to 20.0. A uniform prior with range [0,1] min was used for the time shift parameter *t_s_*. The Hill coefficient *n* and the drug’s elimination half-life—parameters that for biological data could be estimated separately, from a typical PK study—were fixed at *n* = 1 and *t*_1/2_ = 41 minutes. *E_max_* and the coefficients of the signal drift function *a*_0_ + *a*_1*t*_ + *a*_2*t*_^2^ were marginalized.

Since tissues with high values of *EC*_50_ respond less to a given dose of drug, *i.e*. *E* ≪ *E_max_*, the ratio *SD*/*E_max_* ≪ *SD*/*E* underestimates the effect of noise relative to the observed effect. Therefore we computed a signal-to-noise ratio (SNR) to simplify comparisons across the various input values of *EC*_50_ and noise. We defined “signal” as the maximum value of *E*(*C*(*t*)), without added noise, for 0 ≤ *t* ≤ 40min, *i.e*. the local maximum of the modeled signal shortly after the last dose of drug, less the input linear drift at that same time point. In Figure 1 this value can be appreciated near the right side of the plot and ranges from about 3 for *EC*_50_ = 7.5 to about 9 for *EC*_50_ = 0.25. We define SNR as the ratio of this signal to the standard deviation of the added noise.

### Testing the method on *in vivo* data

We tested the model described above using the same phMRI (pharmacological fMRI) data we analyzed previously with the iterative method, namely, regional BOLD-sensitive fMRI time-signal curves from midbrain and striatum in each of two animals (Black et al., 2013). Each animal was studied twice, at least 2 weeks apart, producing 8 regional time-signal curves. On each day a total of 0.1 mg/kg of the dopamine *D*_1_ agonist SKF82958 was given intravenously, divided into 4 equal doses on one day and into 8 equal doses on the other day (Black et al., 2013, Table 4 and Figure 10).

The iterative analysis had allowed only values of 5 or 30 minutes for the half-life of drug disappearance from the blood during the scan session; here we used a uniform prior probability over [2,60] minutes for *t*_1/2_. Prior probabilities for all other parameters were the same as described above for the simulated data.

## Results

### Simulated data

#### Example

Figure 2 provides an example result from one time course, to orient the reader to the following summary. Note that the parameter estimates are (approximately) the best estimates for the provided noisy data, even though they differ slightly from the input values used to produce the data.

**Figure 2.**
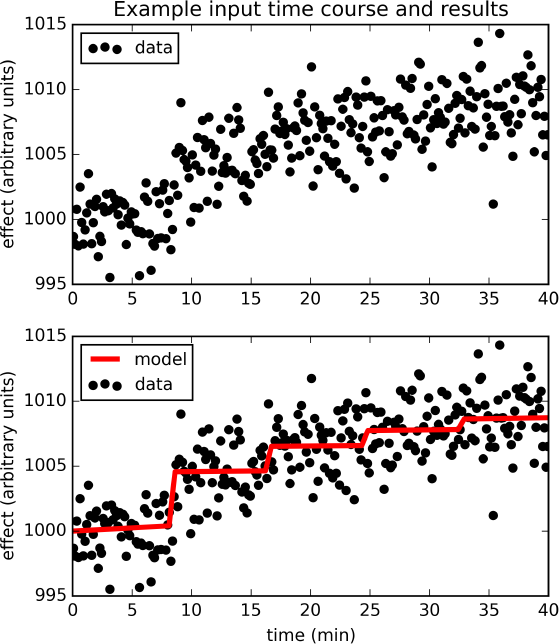
The upper panel shows simulated dose-effect data generated using *E_max_* = 10.0, *EC*_50_ = 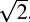, *t_s_* = 0.50, added to 1000+.05*t* +0*t*^2^ and Gaussian noise with SD= 2. In the lower panel, superimposed on the data is the predicted time course of drug effect over time, drawn using the parameter values returned by the Bayesian Data-Analysis Toolbox as most likely given these data and the PKPD model: *E_max_* = 10.6, *EC*_50_ = 1.43, *t_s_* = 0.451, 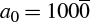, *a*_1_ = 0.0553, *a*_2_ = –0.000149. For this time course, *p*(model) was estimated as 0.540, and the SD of the residuals was 2.04.

#### Sensitivity: p(model) with signal

The full PKPD model explained the data better than a simpler model, *i.e*. *p*(model) >0.5, except when signal was low (higher *EC*_50_) or noise was substantial (Figures 3, 4).

**Figure 3.**
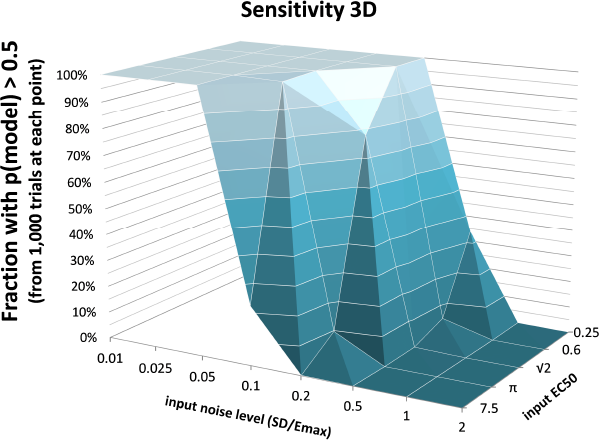
The fraction of time courses for which p(model) >0.5 is shown on the vertical axis as a function of the *EC*_50_ and SD used to generate the time courses.

**Figure 4.**
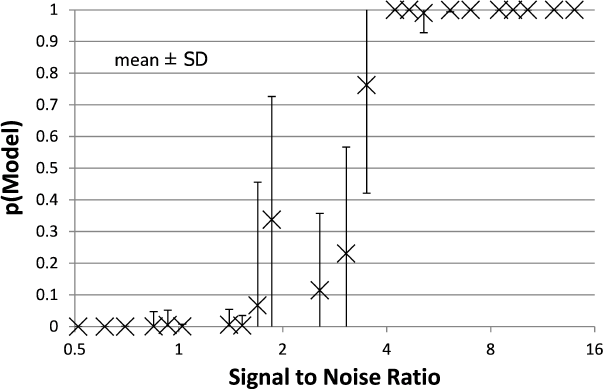
The mean ± SD probability of the full PKPD model is shown for each combination of *EC*_50_ and noise as a function of that combination’s SNR as defined in Methods. Points with SNR outside the range shown here are omitted for clarity.

#### False positives: p(model) with noise only

For the data sets containing no intentional signal, *i.e*. noise added to the *E_max_* = 0 line, the Toolbox never returned *p* > 0.5 for any of the 8,000 curves. In other words, there were no false positives.

#### Accuracy

Accuracy of the *EC*_50_ estimate was considered for time courses with p(model) >0.5. Figure 5 shows the mean estimated *EC*_50_ as a function of the input *EC*_50_; as expected, accuracy is best with higher SNR. Figure 6 shows the ratio of estimated *EC*_50_ to input *EC*_50_ in terms of SNR. Perfect accuracy would produce a ratio of 1.0, and values >1.0 indicate overestimation of *EC*_50_, *i.e*. underestimation of the sensitivity to drug.

**Figure 5.**
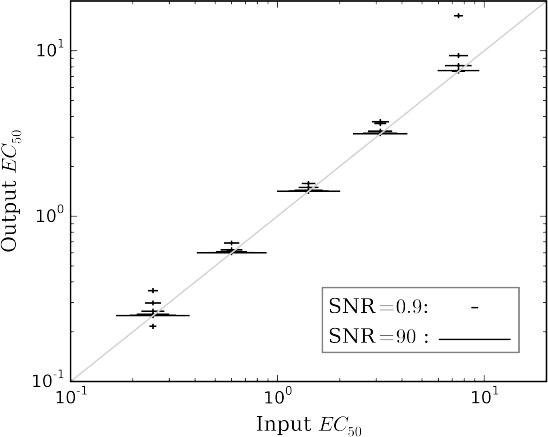
The mean accuracy of the estimated *EC*_50_ for time courses with *p*(model)> 0.5 is shown as a function of the input *EC*_50_. SNR for each estimate is shown by the width of the marker, as indicated by the legend at lower right. The diagonal line indicates equality, *i.e*., perfect accuracy.

**Figure 6.**
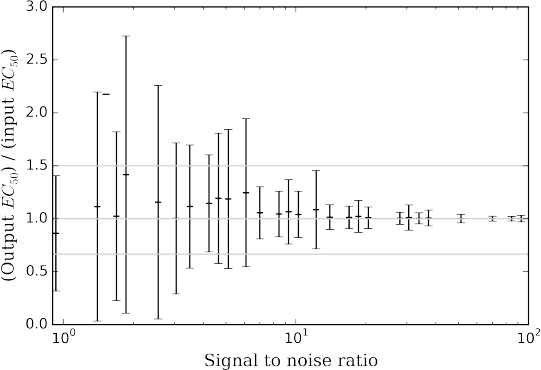
The mean ± SD accuracy of the estimated *EC*_50_ for time courses with *p*(model) >0.5 is shown as a function of SNR as defined in Methods. Here accuracy is defined as the output *EC*_50_ divided by the input *EC*_50_. The full-width horizontal lines indicate perfect accuracy (ratio = 1.0) and 3/2 and 2/3 of perfect accuracy. The accuracy of the estimated *EC*_50_ is superb when SNR > about 6.5, and tends to be accurate for SNR as low as 0.9.

#### In vivo data

The full PKPD model was selected for 6 of the 8 regional time-signal curves (see Table 1). The data and selected model curves are shown in Figure 7.

**Table 1.** Model select results from *in vivo* data

| Panel | Doses | Animal | Region | Prob. | $E_{max}$ | $EC_{50}^*$ | $t_{1/2}$ | $t_s$ |
| --- | --- | --- | --- | --- | --- | --- | --- | --- |
| A | 4 | 1 | Midbrain | 1.00 | 12.59 | 3.44 | 58.33 | 0.98 |
| B | 4 | 1 | Striatum | 1.00 | −13.58 | 4.15 | 59.48 | 0.81 |
| C | 4 | 2 | Midbrain | 1.00 | 29.27 | 6.32 | 3.93 | 0.23 |
| D | 4 | 2 | Striatum | 1.00 | −2.48 | 0.001 | 40.58 | 0.01 |
| E | 8 | 1 | Midbrain | 0.00 | — | — | — | — |
| F | 8 | 1 | Striatum | 0.02 | — | — | — | — |
| G | 8 | 2 | Midbrain | 0.76 | 7.38 | 0.418 | 13.16 | 0.18 |
| H | 8 | 2 | Striatum | 1.00 | −13.9 | 1.63 | 2.00 | 0.72 |

**Figure 7.**
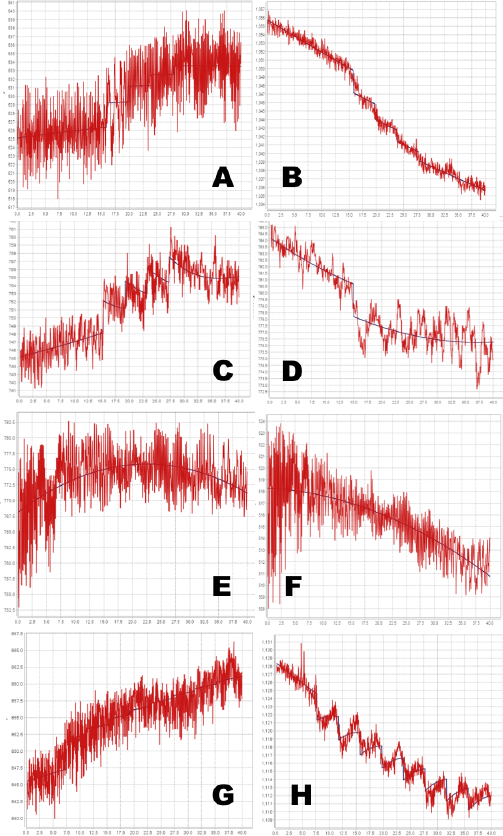
Time-signal curves from *in vivo* data from a phMRI study, in red, with the selected model in dark blue. A-D, 4-dose experiments. E-H, 8-dose experiments. Left column, midbrain. Right column, striatum. See Table 1 for further details.

## Discussion

### Simulation testing

Bayesian parameter estimation for the QuanDyn™ quantitative pharmacodynamic imaging method produced excellent results in simulated data: first, the Model Select method very accurately identified time courses with a meaningful drug-related signal, until noise overwhelmed signal, *i.e*. when SNR < about 3.5. The Bayesian Data-Analysis Toolbox successfully avoided false positives, correctly refraining from identifying a signal in every noise-only time course, even where sensitivity was 100%. In time courses with a signal, mean accuracy was reasonable even in the face of low SNR, as shown in Figures 5 and 6. Furthermore, the errors were conservative, with *EC*_50_ usually erring on the high side (figure 6). Said differently, the most likely quantitative error was to report slightly lower sensitivity to drug, especially when sensitivity is in fact low.

### Limitations

This simulation used a simple noise model that may be best suited to a temporally stable, quantitative outcome measure, such as positron emission tomography, arterial spin labeling, or quantitative BOLD. However, because the PKPD model *E*(*C*) is simply added to the baseline model *B*(*t*), the latter can be replaced with a more complex signal, if needed, for non-quantitative imaging methods. For instance, Fourier series have been used to model typical BOLD-sensitive fMRI data over long time intervals. The baseline model *B*(*t*) could be optimized further to best suit a specific scanner, tracer or sequence, or to other experimental design choices.

Similar comments hold for the signal as well as for noise: the QuanDyn™ quantitative pharmacodynamic imaging method will perform less well if the PKPD model does not realistically model the data. However, prior to initiating an expensive imaging study, one would determine the appropriate family of PKPD models for the drug to be tested, based on traditional dose-response experiments. We discuss this point further in (Black et al., 2013). The choice of imaging method also affects the signal characteristics; for instance, typical BOLD implementations may not provide adequately linear responses to biological signal. On the other hand, using a more traditional phMRI design, the magnitude of the acute BOLD response to a single dose of drug per imaging session did increase monotonically with larger doses (Miller et al., 2013).

### In vivo data

Even with the relatively simple signal and noise models adopted for this initial testing, the tested method appeared to handle reasonably the *in vivo* data from a BOLD phMRI study (Figure 7). Further validation will require a larger set of similar multi-dose phMRI data, and comparison data from a more traditional dose-response study design.

The QuanDyn™ method described here has several potential advantages compared to the traditional approach to quantifying a drug effect, which is to estimate the population *EC*_50_ by sampling a wide range of doses, one dose per subject and several subjects per dose. That approach is an excellent choice when the population under study is homogeneous (*e.g*. an inbred rodent strain), but does not apply well to single human subjects. One might adapt the traditional approach by repeatedly scanning a single subject, one dose per scan session, but that option brings its own complications, including scientific concerns such as sensitization or development of tolerance with repeated doses in addition to the practical and ethical consequences of repeated scanning sessions in each subject. That option, like the population method, would also require that subjects receive doses substantially higher than the *EC*_50_, which may often be inappropriate in early human studies. Specifically, to estimate *EC*_50_, traditional population PKPD studies require drug doses that produce effects of at least ∼ 95%*E_max_* (Dutta et al., 1996). For all these reasons, the QuanDyn™ method may prove to be a better choice when single-subject responses are important, such as for medical diagnosis or individualized treatment dosing. We elsewhere discuss potential challenges related to moving this approach into humans (Black et al., 2013).

## Acknowledgments

Some of these results were presented previously (Koller JM, Bretthorst GL, Black KJ. A novel analysis method for pharmacodynamic imaging. Program #504.1, annual meeting, Society for Neuroscience, Chicago, 20 Oct 2009), and a preprint was posted on bioRxiv (DOI: 10.1101/017921).

## Author Contributions

Jonathan M. Koller performed the experiments, analyzed the data, contributed analysis tools, reviewed and critiqued the manuscript. M. Jonathan Vachon performed the experiments, analyzed the data, reviewed and critiqued the manuscript. G. Larry Bretthorst contributed analysis tools, reviewed and critiqued the manuscript. Kevin J. Black conceived and designed the experiments, performed the experiments, analyzed the data, wrote the paper.

## Data Deposition

The following information was supplied regarding the deposition of related data:

The simulated data sets (1000 time courses for each set of parameter values and noise level) are available at the journal web site as Supplementary Data.

## References

Black, K. J., Koller, J. M., and Miller, B. D. (2013). Rapid quantitative pharmacodynamic imaging by a novel method: Theory, simulation testing and proof of principle. PeerJ, 1:e117.

Bretthorst, G. L. (1988). Bayesian Spectrum Analysis and Parameter Estimation. Lecture Notes in Statistics. Springer-Verlag, New York.

Bretthorst, G. L. (2014). Bayesian data-analysis toolbox, release 4.10, manual version 2. Available at http://bayes.wustl.edu/Manual/BayesManual.pdf (accessed 04 Aug 2015).

Bretthorst, G. L. and Marutyan, K. (2014). Bayesian Data-Analysis Toolbox, v. 4.21 (software). Available at http://bayesiananalysis.wustl.edu (accessed 13 Feb 2015).

Dutta, S., Matsumoto, Y., and Ebling, W. F. (1996). Is it possible to estimate the parameters of the sigmoid *E_max_* model with truncated data typical of clinical studies? Journal of Pharmaceutical Sciences, 85(2):232–239.

Gilks, W. R., Richardson, S., and Spiegelhalter, D. J. (1996). Markov Chain Monte Carlo in Practice. Chapman & Hall/CRC Interdisciplinary Statistics (Book 2). Chapman & Hall, London.

Holford, N. H. G. and Sheiner, L. B. (1982). Kinetics of pharmacologic response. Pharmacology & Therapeutics, 16(2):143–166.

Lavielle, M. (2014). Mixed Effects Models for the Population Approach: Models, Tasks, Methods and Tools. Chapman and Hall/CRC.

Miller, B., Marks, L. A., Koller, J. M., Newman, B. J., Bretthorst, G. L., and Black, K. J. (2013). Prolactin and fMRI response to SKF38393 in the baboon. PeerJ, 1:e195.

